# Sexually dimorphic phenotypes and the role of androgen receptors in UBE3A-dependent autism spectrum disorder

**DOI:** 10.1101/2024.05.02.592248

**Authors:** Yuan Tian, Hui Qiao, Ling-Qiang Zhu, Heng-Ye Man

## Abstract

Autism spectrum disorders (ASDs) are characterized by social, communication, and behavioral challenges. *UBE3A* is one of the most common ASD genes. ASDs display a remarkable sex difference with a 4:1 male to female prevalence ratio; however, the underlying mechanism remains largely unknown. Using the UBE3A-overexpressing mouse model for ASD, we studied sex differences at behavioral, genetic, and molecular levels. We found that male mice with extra copies of *Ube3A* exhibited greater impairments in social interaction, repetitive self-grooming behavior, memory, and pain sensitivity, whereas female mice with UBE3A overexpression displayed greater olfactory defects. Social communication was impaired in both sexes, with males making more calls and females preferring complex syllables. At the molecular level, androgen receptor (AR) levels were reduced in both sexes due to enhanced degradation mediated by UBE3A. However, AR reduction significantly dysregulated AR target genes only in male, not female, UBE3A-overexpressing mice. Importantly, restoring AR levels in the brain effectively normalized the expression of AR target genes, and rescued the deficits in social preference, grooming behavior, and memory in male UBE3A-overexpressing mice, without affecting females. These findings suggest that AR and its signaling cascade play an essential role in mediating the sexually dimorphic changes in UBE3A-dependent ASD.

## Introduction

Autism Spectrum Disorder (ASD) is a neurodevelopmental disorder characterized by impairments in sociability and communication, as well as the presence of restricted and repetitive behaviors ^1^. According to reports from the Centers for Disease Control and Prevention (CDC) of the USA, the prevalence of ASD has been increasing over the past decades. The global prevalence of the disease is estimated at 1%, and approximately 1 in 36 children have been diagnosed with ASD in the United States ^2^. However, the cause of ASD remains less clear despite the identification of various risk factors. Environmental risk factors include parental age, maternal pregnancy, and pregnancy complications ^3^. Population screening studies have shown that genetics play a significant role in the development of the disease, supported by a heritability estimated at 90% in ASD ^4^. Recent research has discovered hundreds of genetic variations, both common and rare, that contribute to the development of ASD ^5^.

Remarkably, ASD has a four-fold higher occurrence in males than in females ^6^. To this day, the exact mechanisms and genetic determinants underlying the sex-bias in autism are still largely unknown. One possibility is related to male-biased risk factors, such as the influence of sex steroid hormone signaling and prenatal programming on brain development, while another possibility pertains to a female-biased protective mechanism that results in a higher mutation “burden” for ASD risk in females. However, the etiology of these possibilities is not well-understood ^7–9^. In addition, different phenotypic presentations of ASD in males and females have been reported in human patients. Males with ASD tend to exhibit more externalizing behaviors such as aggression, hyperactivity, reduced social behavior, and increased restricted and repetitive behaviors ^7, 9–12^, whereas females with ASD show greater internalizing symptoms such as anxiety, depression, and other emotional problems ^7, 12, 13^. Interestingly, sex chromosomes, sex hormone signaling pathways, and sexually dimorphic factors such as differential gene expression in neurons and glial cells, have all been proposed as potential causes of these sex differences ^7, 9^. Despite these possibilities, however, the exact mechanisms underlying sexually dimorphic expression of autistic phenotypes remain unclear.

*UBE3A* is a significant ASD risk gene located on chromosome 15q11-13. The protein product of *UBE3A* functions as a HECT family E3 ligase regulating protein ubiquitination and degradation, as well as a transcription coactivator regulating gene expression ^14^. The *UBE3A* gene is imprinted in the brain, with only the maternal copy of the gene being expressed in neurons, whereas the paternal copy is silenced ^14^. The *UBE3A* gene plays essential roles in brain development, and it is well established that deletions of the *UBE3A* gene result in Angelman syndrome ^15^. In contrast, excessive UBE3A expression and activity resulting from duplication or triplication of maternal chromosome 15q11-13 ^14, 16^, or de novo hyperactivating missense mutations of UBE3A ^17–19^, are implicated in the development of ASD. The 15q11.2 microduplication encompassing only the *UBE3A* gene has been reported to cause learning disability and autistic features in human patients, highlighting the significance of the *UBE3A* gene in ASD development ^20^. ASD diagnoses due to genetic alterations in *UBE3A* account for 1-2% of all autism cases, making it one of the strongest genetic risk factors for autism ^21, 22^.

*Ube3A* 2x transgenic (Tg) mice carry a tripled *Ube3A* gene dosage that mimics the disease condition, and exhibit typical autistic-like behaviors such as impaired social behavior and communication, as well as increased repetitive behavior ^23, 24^. Using the *Ube3A* 2xTg ASD mice, we examined whether sexually dimorphic phenotypes manifest in Ube3A-dependent ASD and investigated the potential mechanisms that underlie the observed sex differences in this disorder. To summarize, we observed male-biased behavioral abnormalities in social interaction, repetitive self-grooming behavior, memory function, and pain sensitivity, which were associated with sexually dimorphic changes in the androgen receptor (AR) signaling pathway. We found that AR was decreased in 2xTg male mice due to enhanced degradation, and more importantly, restoration of AR expression in the brain was able to rescue a large portion of the behavioral deficits in male 2xTg mice.

## Results

### Sexually dimorphic behavioral impairments in Ube3A 2xTg ASD mice

To investigate whether sexually dimorphic changes are present in UBE3A-dependent ASD, we examined the three core autism-relevant behaviors, including impaired social interaction, abnormal social communication, and the presence of restricted and repetitive behaviors in male and female *Ube3A* 2xTg mice. To examine their social behaviors, we used the three-chamber social test, in which a stranger mouse was placed into a cage at either side chamber and a test mouse of the same sex was allowed to move freely in the apparatus to examine its social preference (Fig. 1A). The wild type (WT) male and female mice displayed normal preference to interact with the stranger mice, whereas the 2xTg male mice lost this preference and spent a similar amount of time interacting with the two cages (Fig. 1A, 1C). Surprisingly, female Tg mice preserved the preference to the stranger mouse (Fig. 1A, 1C), indicating social disruptions in Tg male, but not Tg female, mice. We then introduced a second mouse of the same sex (novel mouse) into a cage on the remaining side chamber, and the test mouse was allowed to interact with both mice (Fig. 1B). Quantification of the interaction time in WT animals showed a normal preference to the novel mouse as expected, but both male and female Tg mice spent a similar amount of time interacting with the two conspecifics, indicating that social novelty was impaired in Tg animals regardless of sex (Fig. 1B, 1D). Furthermore, to assess repetitive self-grooming behavior, each test mouse was placed individually into a regular housing cage, and the amount of time spent grooming was quantified. Elevated grooming time was observed in male but not female *Ube3A* 2xTg mice, suggesting that male Tg mice are preferentially affected in their repetitive grooming behaviors (Fig. 1E).

**Fig. 1:**
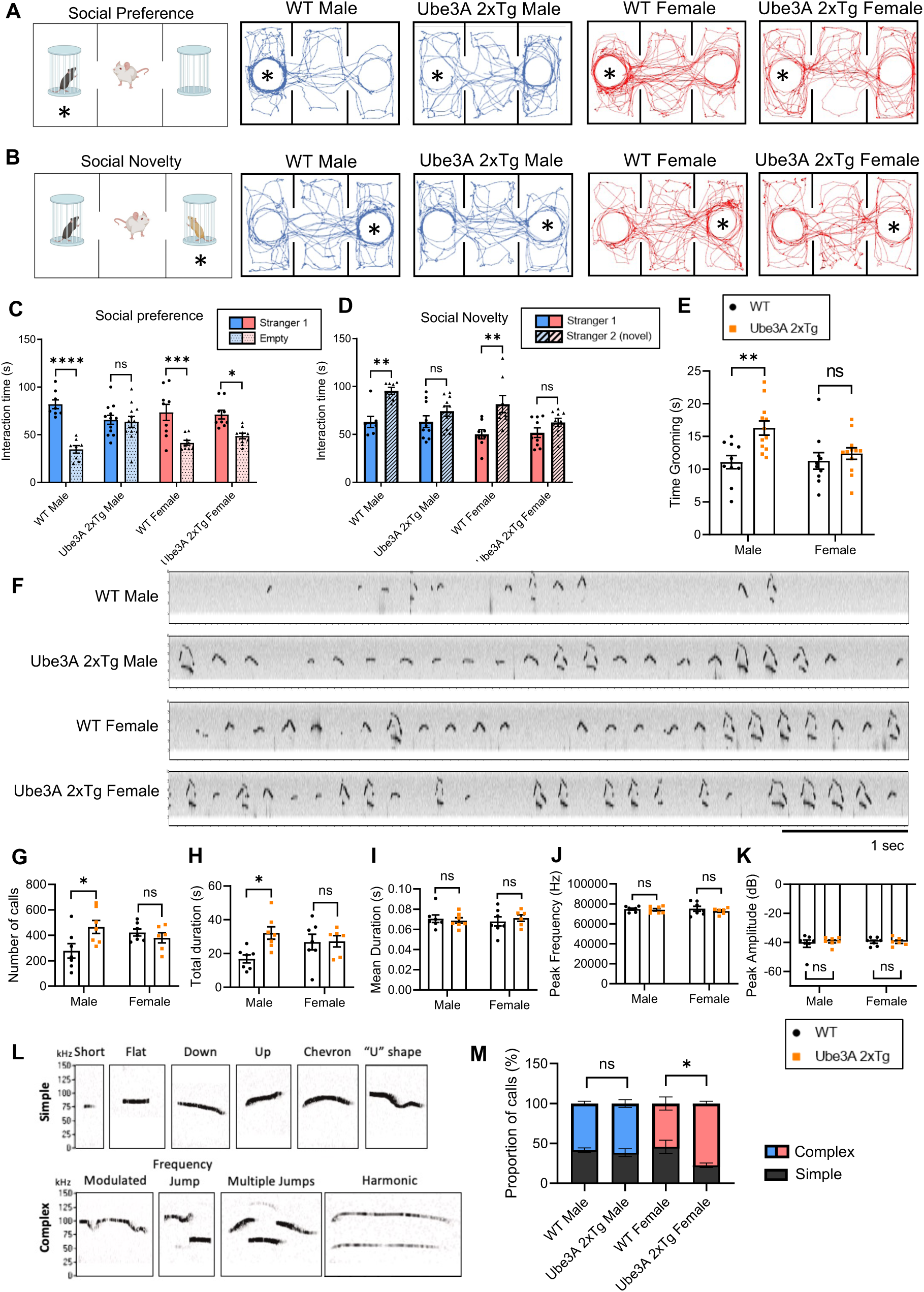
Sexually dimorphic deficits in social interaction, repetitive self-grooming behaviors, and ultrasonic vocalizations in male and female *Ube3A* 2xTg mice. **(A,B)** The paradigm for the three-chamber social test and traces of track paths. For social preference (A), an unfamiliar mouse was placed into either of the side chambers and the test mouse was allowed to move freely in the apparatus. For social novelty (B), a second mouse (novel mouse) was placed into the remaining empty chamber, and the test mouse was allowed to interact with both mice. **(C,D)** Quantification of the interaction time showed a decrease in preference for the stranger mouse (C) and novel mouse (D) in male *Ube3A* 2xTg mice, while female *Ube3A* 2xTg mice showed mild change in social preference (C) but clear impairment in social novelty (D). **(E)** Male but not female *Ube3A* 2xTg mice showed increased repetitive self-grooming behavior. **(F)** Representative vocalization recordings from P5 WT and *Ube3A* 2xTg mice. **(G-K)** Quantification of the number of calls (G), total call duration (H), mean call syllable duration (I), peak frequency (J), and peak amplitude (K) for P5 WT and *Ube3A* 2xTg mice. **(L)** Representative calls of each type used in syllable characterization. **(M)** *Ube3A* 2xTg female animals showed an increase in Complex type calls and a decrease in Simple type calls at P5, while *Ube3A* 2xTg males are similar to WT males. **In (C),** n=8 WT male; n=12 Tg male; n=9 WT female; n=9 Tg female. **In (D),** n=7 WT male; n=10 Tg male; n=8 WT female; n=9 Tg female. **In (E),** n=10 WT male; n=12 Tg male; n=10 WT female; n=12 Tg female. **In (G-M)** n=7 WT male; n=7 Tg male; n=7 WT female; n=6 Tg female. Mean ± SEM. *p<0.05; **p<0.01; ***p<0.001; ****p<0.0001. ns, not significant. In (C), (D), (M) Three-way ANOVA with Bonferroni’s multiple comparisons test; In (E), (G-K) Two-way ANOVA with Bonferroni’s multiple comparisons test.

Impairments in social communication have been established as one of the three main ASD hallmarks. To examine the presence of communication deficits in male and female *Ube3A* 2xTg animals, we recorded and analyzed their ultrasonic vocalizations (USVs). Individual pups isolated from the mother and littermates at P5, P7 and P9 were placed in a recording chamber for 5 min in which a microphone was present to record vocalization signals (Fig. 1F). At P5, but not at P7 and P9, we observed an increase in the number of calls and the total call duration in *Ube3A* 2xTg males compared to WT males (Fig. 1G, 1H, S4A, S4B), a trend also noted in other ASD models ^25–29^. Conversely, no change was observed in the number of calls and total call duration in *Ube3A* 2xTg females at P5, P7 and P9 (Fig. 1G, 1H, S4A, S4B). Furthermore, both male and female Tg animals displayed a normal mean call duration, peak frequency of calls, and peak amplitude of calls at P5, P7 and P9 (Fig. 1I-1K, S4C-E). Further analysis of the call syllable types (Fig. 1L) revealed that at P5, female *Ube3A* 2xTg animals made more complex USV calls than WT females (Fig. 1M), characterized by a significant increase in harmonic calls and a decrease in chevron calls (Fig. S1B). In contrast, male Tg mice used call syllable types similar to WT males at P5 (Fig. 1M, S1A). At P7 and P9, both male and female Tg mice exhibited call syllable types comparable to those of their WT counterparts (Fig. S4F, S4G). These findings suggest that UBE3A over-dosage results in altered communication in both male and female animals at P5, but in different manners, with male Tg mice increasing their number of calls and total call duration, while female Tg mice using more complex syllables.

Intellectual and cognitive abnormalities are one of the most common comorbidities seen in human ASD ^30, 31^. We therefore sought to examine whether memory deficits are differentially affected in male and female *Ube3A* 2xTg animals using the Novel Object Recognition (NOR) test and Barnes maze test. For the NOR test, two identical objects were introduced to the test mouse, and 4 hours or 24 hours later, one of the familiarized objects was replaced with a novel object (Fig. 2A). The Tg animals showed normal short-term memory 4 h after training, regardless of sex (Fig. 2B). However, while Tg females displayed intact recognition capabilities after 24 h, Tg males failed to recognize the familiar object, indicating that long-term memory was impaired specifically in *Ube3A* 2xTg males (Fig. 2C). In the Barnes maze test for spatial memory, mice were trained over a period of 4 days to learn the location of a target escape hole on a circular board containing 20 holes along its circumference, with spatial cues placed around the testing room to help animals navigate and locate the target hole (Fig. 2D, 2E). 24 hours and 5 days after training sessions, the test mouse was returned to the board with the target hole covered, and the number of errors made before reaching the escape hole was quantified (Fig. 2D, 2E). In the 24 h probe test, neither male nor female Tg animals showed significant differences in the number of errors as compared to WT mice (Fig. 2F). However, probing long-term memory retention 5 d after training showed that male, but not female, Tg mice made significantly more primary errors than their WT counterparts (Fig. 2G), Together with the results from the NOR test, these findings indicate that *Ube3A* 2xTg males are preferentially affected in long-term memory in contrast to *Ube3A* 2xTg females.

**Fig. 2:**
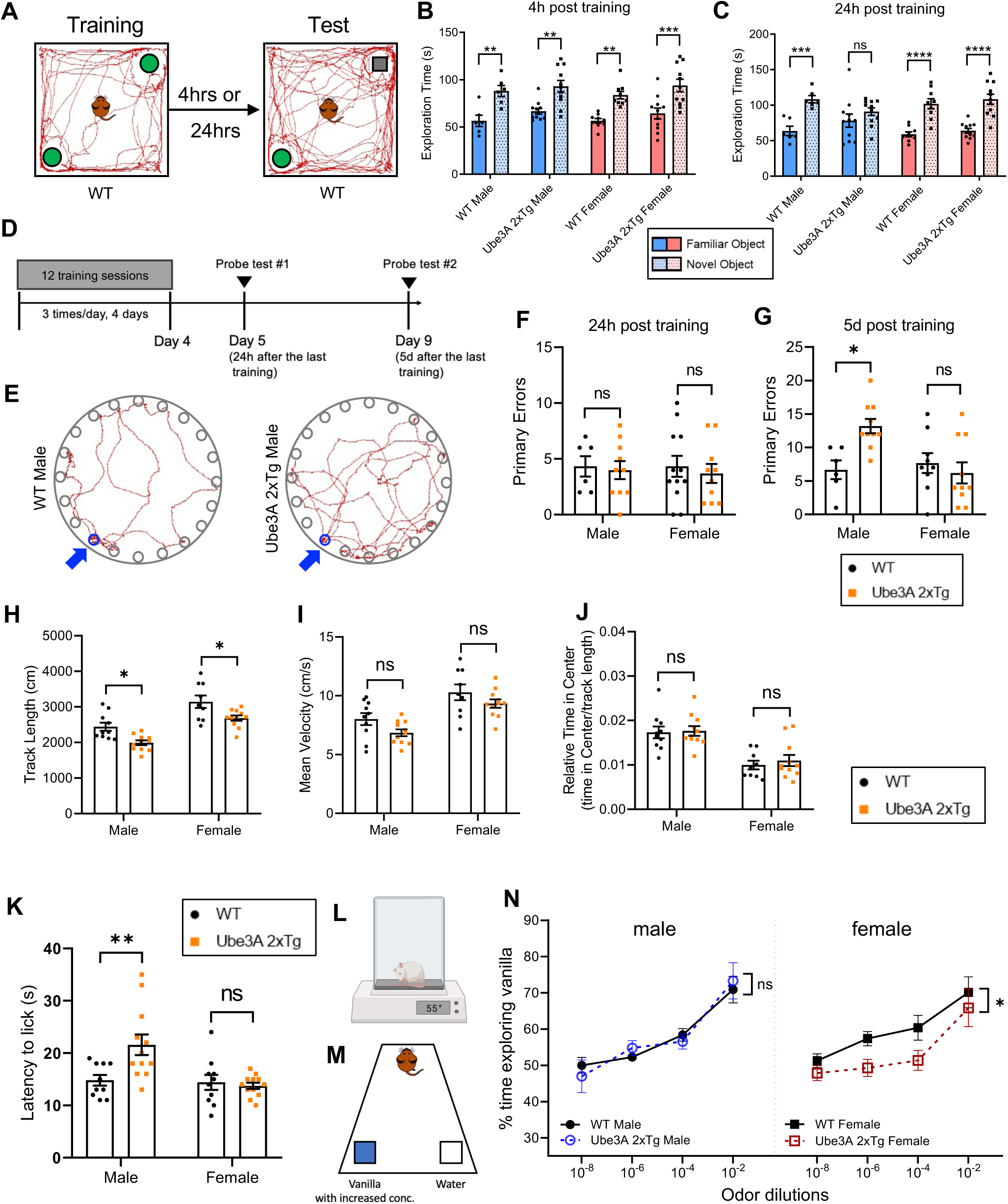
Sexually dimorphic deficits in memory, sensory functions, but not locomotion in male and female *Ube3A* 2xTg mice. **(A)** The paradigm for novel object recognition test. **(B,C)** Exploration time of familiar object vs. novel object 4 h (B) and 24 h (C) post training. *Ube3A* 2xTg animals displayed intact short time memory (B), and only Tg males showed impaired long-term memory (C). **(D)** The timeline of Barnes spatial memory maze. **(E)** Traces of track paths during the 5 d memory probe for WT and *Ube3A* 2xTg males. Blue arrows point to the escape hole. **(F,G)** Primary errors made to find the escape hole 24 h (F) and 5 d (G) after the last training. *Ube3A* 2xTg animals displayed intact short time memory (F), and only Tg males showed impaired long-term memory (G). **(H-J)** Both male and female *Ube3A* 2xTg mice showed decreased track lengths (H), normal mean velocities (I) and no change in relative time in center (J) in the open-field test. **(K,L)** Test mice were put on 55℃ hot plate and their latency to lick a hindpaw was recorded. Male *Ube3A* 2xTg mice showed reduced pain sensitivity compared to WT males, while Tg females were intact. **(M)** The paradigm for the Olfactory Detection Threshold Test. Test mice were exposed to increased concentration of vanilla. Time spent exploring vanilla vs. water was recorded. **(N)** Male *Ube3A* 2xTg mice showed intact olfactory sensitivity compared to WT males, whereas Tg females showed reduced olfactory sensitivity compared to WT females. **In (B-C),** n=6 WT male; n=11 Tg male; n=9 WT female; n=11 Tg female. **In (F)**, n=6 WT male; n=10 Tg male; n=12 WT female; n=10 Tg female. **In (G)**, n=6 WT male; n=10 Tg male; n=9 WT female; n=10 Tg female. **In (H-J)**, n=10 WT male; n=11 Tg male; n=9 WT female; n=11 Tg female. **In (K)**, n=10 WT male; n=12 Tg male; n=10 WT female; n=12 Tg female. **In (N)**, n=10 WT male; n=10 Tg male; n=9 WT female; n=10 Tg female. Mean ± SEM. *p<0.05; **p<0.01; ***p<0.001; ****p<0.0001. ns, not significant. In (B), (C), (N) Three-way ANOVA with Bonferroni’s multiple comparisons test; In (F-K) Two-way ANOVA with Bonferroni’s multiple comparisons test.

Next, in the Open-field test, although the 2xTg animals displayed normal locomotion, both sexes showed a significant decrease in their total track length (Fig. 2H). Quantification of the average velocity and the time spent in the center of the open field normalized to the total distance traveled showed no difference between WT and Tg mice within each sex (Fig. 2I, 2J). However, in both genotypes, female mice generally exhibited a longer track length, higher mean velocity, and spent less relative time in the center compared to male mice (Fig. 2H-2J). Together, these results indicate that *Ube3A* 2xTg animals are less active in general, without signs of enhanced anxiety.

In addition, atypical sensory functioning has often been found in ASD patients and animal models, such as impaired peripheral somatosensory functions ^32–34^ and odor perception ^35, 36^. We first performed the hot plate assay to evaluate the pain sensitivity of *Ube3A* 2xTg mice by measuring the latency to lick their hind paw after being placed on a surface heated to 55 °C (Fig. 2L). Male Tg mice displayed reduced sensitivity to the hot temperature, whereas female Tg had intact pain sensitivity (Fig. 2K). We then examined their olfactory sensitivity using the olfactory detection threshold test, in which the mice were exposed to increased concentrations of vanilla, and the time spent exploring the vanilla odor versus the water control was recorded (Fig. 2M). We observed normal detection of the vanilla odor in 2xTg male mice, but the female Tg animals were unable to perceive the odor until the highest concentration was applied (Fig. 2N), indicating a reduced olfactory sensitivity specific to female, but not male, 2xTg animals.

### The androgen receptor signaling pathway is dysregulated mainly in male Ube3A 2xTg ASD mice

Sex hormones act throughout the entire brain of both males and females, regulating many neural and behavioral functions ^37–40^. Sex hormones bind to and activate sex hormone steroid receptors, either androgen receptors (AR) or estrogen receptors (ER), and the complexes will then translocate into the nucleus to regulate gene expression ^37, 41^. Importantly, the sex differences observed across several brain functions are mediated by sex hormone signaling pathways ^37^. Both AR and ER are known to be regulated by E3 ubiquitin ligases, which target them for ubiquitination and degradation ^42–45^. Given that UBE3A functions as an E3 ligase, it is intriguing to consider whether UBE3A regulates the turnover of AR and ER, which may contribute to the sexually dimorphic changes in *Ube3A* 2xTg animals. To this end, we examined the expression of AR, and ER alpha (ERα) and beta (ERβ) in prefrontal cortical lysates of 2xTg brains by western blotting. Surprisingly, quantification analysis revealed decreased expression of AR in both male and female *Ube3A* 2xTg animals, whereas ERα and ERβ expression were largely intact compared to their WT counterparts, regardless of sex (Fig. 3A, 3B). Consistently, by immunostaining of AR in cortical brain slices, we observed a reduction of AR immunointensity in both male and female 2xTg brains (Fig. 3C, 3D). Given that at basal conditions, AR activity is significantly lower in female brains due to low androgen hormone levels compared to males, we predicted that AR reduction should affect downstream gene expression mainly in males ^46, 47^. To test the idea, we performed RNAseq assays using 2xTg mouse brain tissues and analyzed the androgen-regulated genes (ARGs) and estrogen-regulated genes (ERGs) in comparison with their WT counterparts. As expected, ARGs were found to be markedly downregulated in male *Ube3A* 2xTg brains, but not in female 2xTg brains (Fig. 3E), indicating a minimal downstream disturbance caused by AR reduction in female 2xTg brains. On the other hand, little to no ERGs were dysregulated in Tg animals, regardless of sex (Fig. 3F). These findings suggest that the AR signaling pathway is disrupted in *Ube3A* 2xTg animals, which leads to strong genetic disturbances specifically in males.

**Fig. 3:**
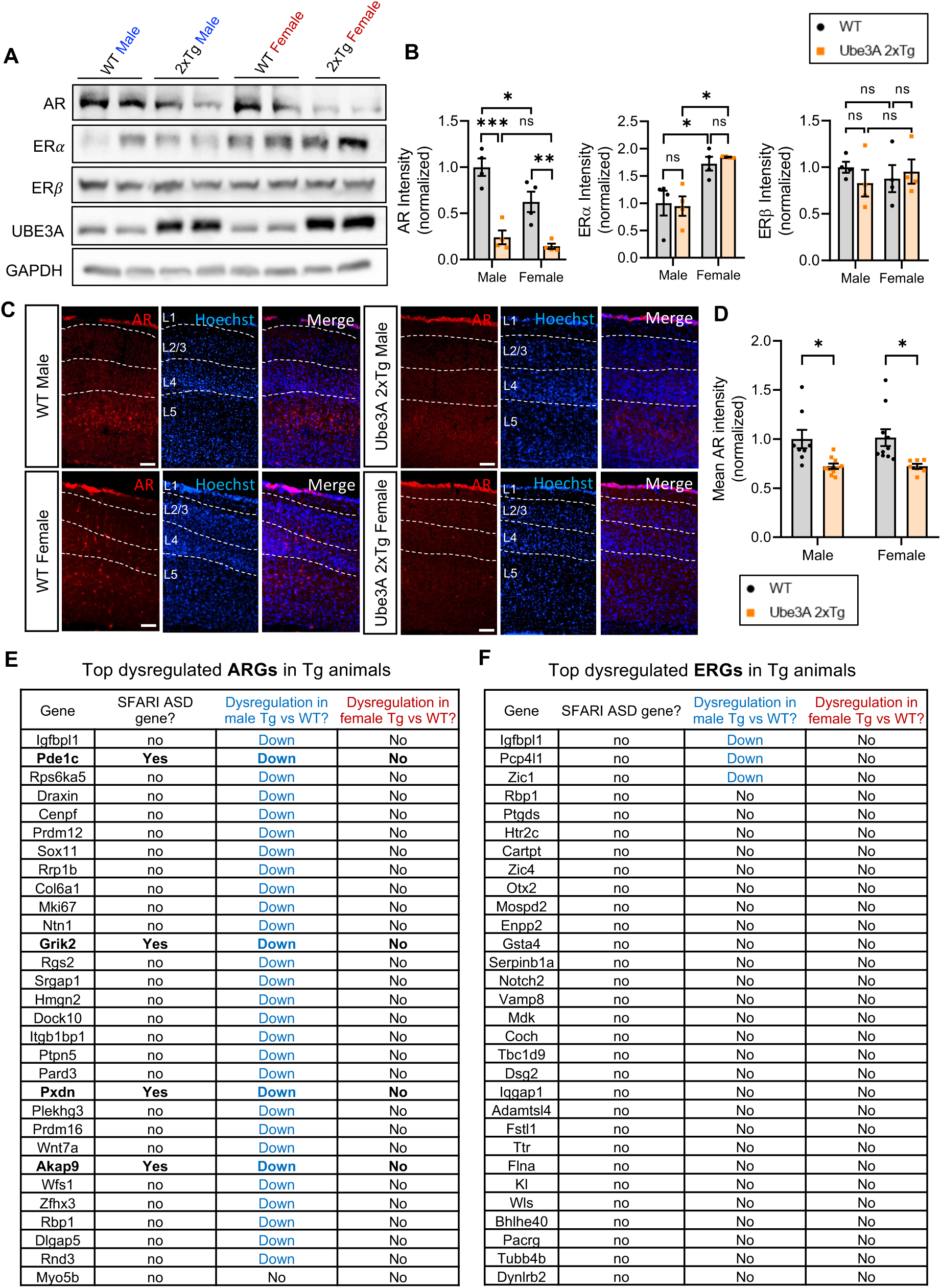
Androgen receptor expression is reduced in *Ube3A* 2xTg animals, leading to dysregulation of androgen responsive genes specifically in males. **(A,B)** Androgen receptors (AR) were reduced in both Tg male and Tg female prefrontal cortical lysates at P7. Estrogen receptors alpha (ERα) and beta (ERβ) didn’t change in *Ube3A* 2xTg animals compared to WT counterparts. n=4 animals/group. **(C,D)** Immunohistochemistry of WT and Tg cortical slices at P60 showed a marked decrease in AR immunointensity in Tg animals. n = 8 WT male; n = 10 Tg male; n = 10 WT female; n = 7 Tg female. Scale bars, 100 μm. **(E,F)** Top dysregulated androgen responsive genes (ARGs, E) and estrogen responsive genes (ERGs, F) in *Ube3A* 2xTg male and female brains, compared to their WT counterparts (n=3 animals/group). Blue fonts indicate dysregulation in male *Ube3A* 2xTg brains compared to WT males. Red fonts indicate dysregulation in female *Ube3A* 2xTg brains compared to WT females. Bold fonts indicate known SFARI ASD genes. ARGs were strongly dysregulated in Tg males but not Tg females. Little to no ERGs were dysregulated in Tg animals. Mean ± SEM. *p<0.05; **p<0.01; ***p<0.001. ns, not significant. Two-way ANOVA with Bonferroni’s multiple comparisons test.

### Reduction of the androgen receptor in Tg mice is caused by UBE3A-mediated ubiquitination and degradation

To verify whether AR reduction is dependent on an elevated dosage of UBE3A, we transfected primary cortical neurons with UBE3A and analyzed AR expression via immunocytochemistry (ICC) at DIV 8 (Fig. 4A). We found that the AR level decreased significantly in neurons overexpressing UBE3A as compared to the control (Fig. 4B), indicating that UBE3A overexpression causes AR reduction.

**Fig. 4:**
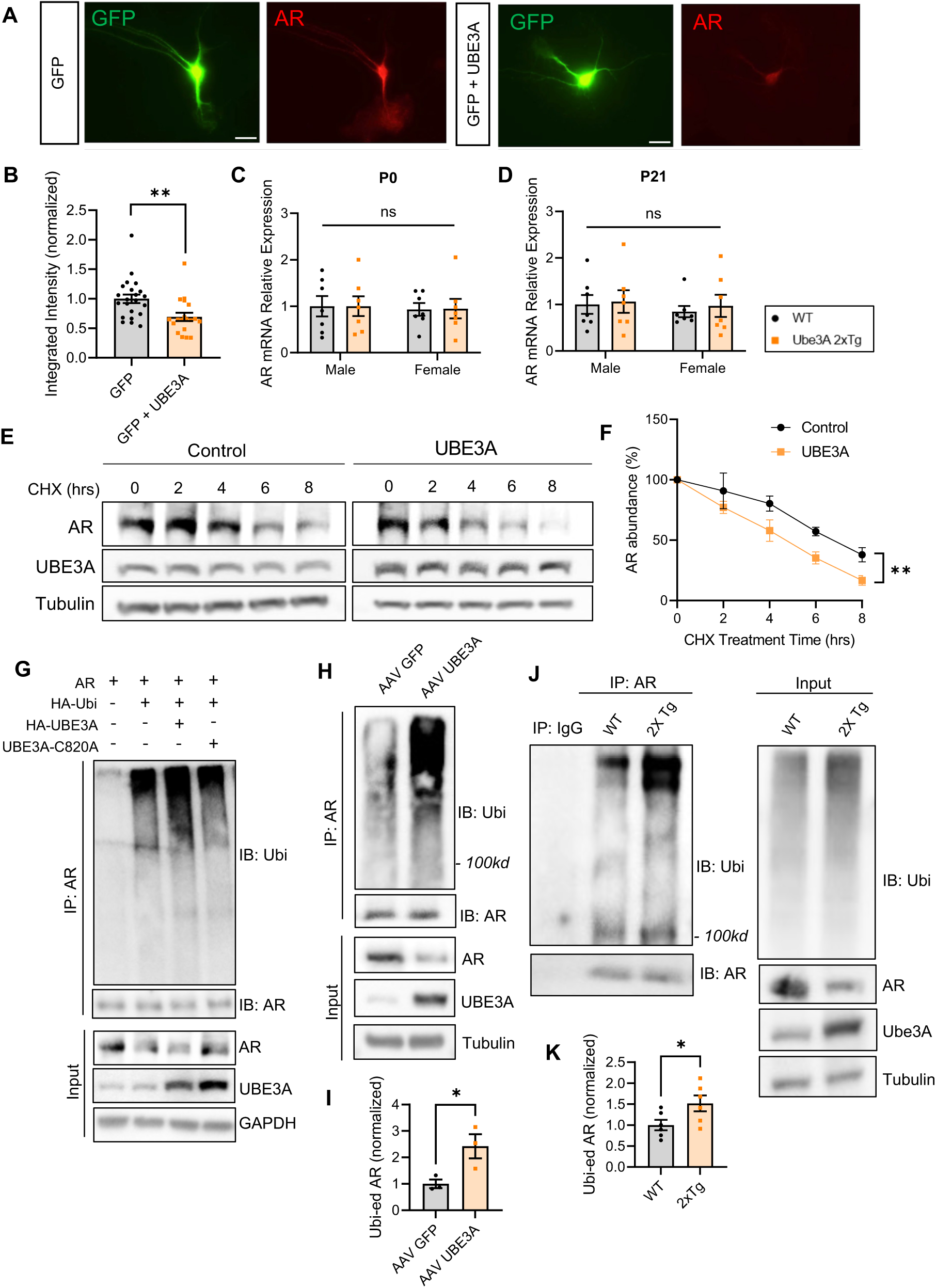
UBE3A overexpression can independently induce AR reduction by increasing ubiquitination and degradation of AR. **(A)** Representative images of immunocytochemistry for AR (red) at DIV 8 in rat cortical neuron cultures after transfected with GFP alone or together with UBE3A at DIV4. Scale bars, 25 μm. **(B)** UBE3A overexpression reduced AR intensity in primary culture neurons. GFP: n=22, GFP+UBE3A: n=19. **(C,D)** AR mRNA expression detected by qPCR was not altered in Tg males and Tg females at postnatal day 0 (C) and postnatal day 21 (D). P0: n=7 animals/group; P21: n=7 animals/group. **(E)** Degradation assay of AR with or without UBE3A. Transfected HEK cells were treated with cycloheximide (CHX) for various time points and cell lysates were collected to examine AR levels by Western blot. **(F)** Quantification of the degradation rate of AR over time; n=3 independent experiments. **(G)** AR ubiquitination assay. HEK293T cells were transfected with AR, HA-ubiquitin (Ubi), and either a vector control, UBE3A, or the E3 ligase dead mutant UBE3A C820A for 2 d. AR was immunoprecipitated and probed for ubiquitin. Cell lysates (input) were also probed to detect total protein levels. **(H,I)** AR ubiquitination assays using lysates of neurons infected with AAV2 GFP or AAV2 UBE3A virus for 10 d (H). Increased intensity of ubiquitination signals on AR was detected (I). n=3 independent experiments. **(J,K)** AR was immunoprecipitated from brain lysates and probed for ubiquitin signals. An elevated level in AR ubiquitination was detected in *Ube3A* 2xTg mice compared to WT. n=6 animals/group. Mean ± SEM. *p<0.05; **p<0.01. ns, not significant. In (C), (D), (F) Two-way ANOVA with Bonferroni’s multiple comparisons test; In (B), (I), (K) Unpaired two-tailed t test.

The UBE3A-induced reduction in AR could result from facilitated degradation or altered transcription. To clarify this, we performed real-time PCR (qPCR) on WT and Tg brain lysates. At either postnatal day 0 or postnatal day 21, no significant changes in AR mRNA level were observed in Tg animals, regardless of sex (Fig. 4C, 4D).

To determine whether the degradation of AR is altered by UBE3A, we transfected HEK cells with AR alone or together with UBE3A for 48 h. Cells were then incubated with the protein translation inhibitor cycloheximide for 0, 2, 4, 6, 8 h before being harvested for western blot analysis. We found that while the AR levels decreased over time due to degradation, the rate of decay became faster in cells co-transfected with UBE3A (Fig. 4E, 4F), indicating that UBE3A expression enhanced the degradation of AR.

Given that UBE3A is a ubiquitin protein ligase, the change in AR turnover by UBE3A could result from changes in its ubiquitination status. To test this possibility, we transfected HEK cells with AR, HA-Ubi, together with either HA-UBE3A, or the ligase dead mutant UBE3A C820A, for 2 d. AR was immunoprecipitated and probed for ubiquitin. Indeed, the ubiquitin signals of AR were significantly increased in cells co-transfected with UBE3A as compared to the control (Fig. 4G). In contrast, no changes in AR ubiquitination were observed in cells co-transfected with the ligase dead mutant UBE3A C820A (Fig. 4G). Moreover, a significant decrease in AR was detected in UBE3A-overexpressing cells (Fig. 4G). To further confirm the role of UBE3A on AR ubiquitination, we performed a ubiquitination assay in neurons infected with an AAV2 virus of either GFP or UBE3A for 10 d. Indeed, compared with GFP, the UBE3A virus caused a significant increase in AR ubiquitination (Fig. 4H, 4I). In addition, ubiquitination assays using lysates from mouse brain tissues also showed increased AR ubiquitination in UBE3A over-dosage brains as compared to WT brains (Fig. 4J, 4K). These results suggest that AR is targeted by UBE3A for ubiquitination and degradation.

### AR restoration in Ube3A 2xTg brains normalizes the expression of ASD-associated ARGs

Following binding with androgens, the activated ARs will translocate into the nucleus to regulate gene expression. In addition to mediating sex hormone signaling and participating in important brain functions ^37, 39, 40, 46, 48^, *AR* and many AR-regulated genes are known to be associated with ASD ^5, 49–61^. Indeed, *AR* itself and at least seven ARGs are listed in the SFARI ASD gene database ^5^. Additionally, studies have shown that dysregulation of AR is associated with the development of several autistic features, including altered social behaviors ^51, 62^, impaired spatial learning and memory ^63, 64^, and hyperactivity symptoms such as attention deficit hyperactivity disorder (ADHD) ^52, 65^. Therefore, the AR system may play an important role in mediating the sexually dimorphic alterations in *Ube3A* 2xTg ASD mice.

To test this possibility, we performed bilateral intracerebroventricular (ICV) injections of a GFP-AR virus (LV-GFP-AR) at postnatal day 0 to replenish AR in *Ube3A* 2xTg brains, or a GFP virus (LV-GFP) as a control (2 μl per side) in WT and Tg pups (Fig. 5A). Animals were later subjected to different behavioral tests at postnatal day 30 to 55. The infected animal brains were processed for biochemical analysis at postnatal day 30 to 60. Imaging of brain slices confirmed the expression of GFP-AR and GFP control virus in the prefrontal cortex (PFC) and hippocampus (HPC). Immunohistochemistry (IHC) of AR also showed co-localization of elevated AR immunosignals with GFP-AR virus-infected green cells in the PFC and HPC (Fig. 5A).

**Fig. 5:**
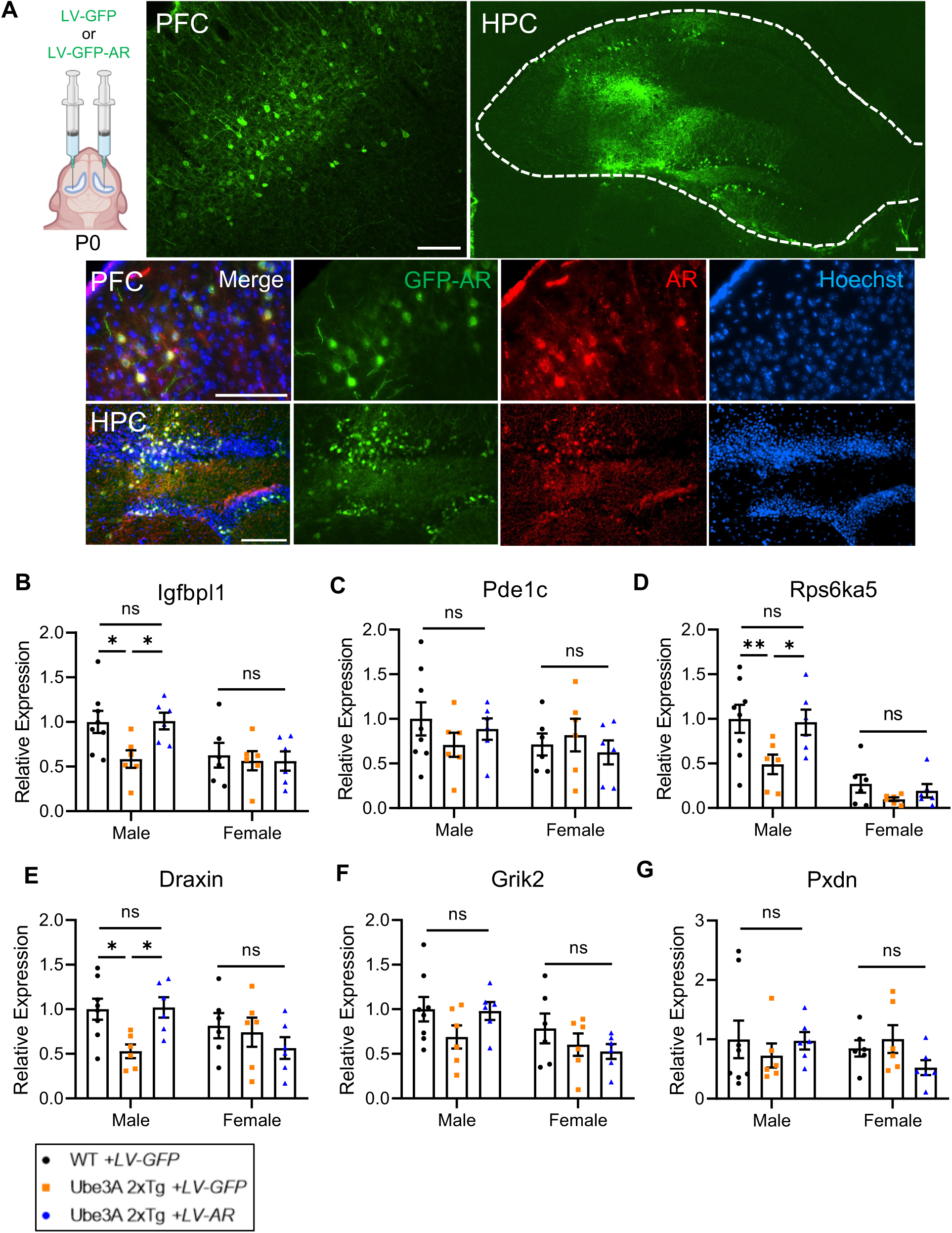
AR restoration in *Ube3A* 2xTg mouse brains rescues expression of androgen responsive genes in males, without affecting females. **(A)** Intracerebroventricular injection of GFP (LV-GFP) or GFP-fused AR (LV-GFP-AR) lentivirus at postnatal day 0. The mice were perfused at postnatal day 30 to 60 for cryostat to validify viral expression. LV-GFP-AR infection significantly increased AR expression in PFC and HPC. PFC, prefrontal cortex. HPC, hippocampus. Scale bars, 100 μm. **(B-G)** Expressions of six ASD-associated androgen responsive genes were rescued in *Ube3A* 2xTg males by ICV injection of LV-GFP-AR at postnatal day 0, detected by qPCR in postnatal day 40 prefrontal cortical lysates. WT male+LV-GFP: n=8, Tg male+LV-GFP: n=6, Tg male+LV-GFP-AR: n=6, WT female+LV-GFP: n=6, Tg female+LV-GFP: n=6, Tg female+LV-GFP-AR: n=6. Mean ± SEM. *p<0.05; **p<0.01; ns, not significant. Two-way ANOVA with Bonferroni’s multiple comparisons test.

We next wanted to determine whether AR restoration could rescue the dysregulation of ARGs in *Ube3A* 2xTg males. To test this idea, we selected 6 candidates out of the top dysregulated ARGs in male *Ube3A* 2xTg brains (Fig. 3E) which are known to be associated with ASD ^5, 53–61^, and examined whether their expression was rescued by LV-GFP-AR.

Of the 6 ARG candidates, dysregulated candidate *lgfbpl1* encodes insulin-like growth factor-binding protein like 1, which acts as a PI3K-AKT pathway inhibitor ^66^. Upregulation of this pathway is involved in the pathogenesis of autism, and has become a common symptom in human patients as well as a focal point in autism studies ^53, 54, 67^. Inherited missense variants and intronic single nucleotide polymorphisms (SNP) within the second candidate gene, *Pde1c*, have been associated with ASD ^55–57^. The *Rps6ka5* gene is located on 14q32.11, and its microdeletion has been found to cause developmental and language delay in human patients ^58^. *Draxin* is involved in the regulation of axon extension, and several human neurological disorders including intellectual disability and ASD are known to be associated with aberrant axon development ^59^. The last two dysregulated candidate ARGs are *Grik2*, whose mutations have also been associated with ASD ^60, 68, 69^, and *Pxdn*, duplication of which has been observed in human autism patients ^61^.

Real-time PCR quantification of mRNA from prefrontal cortical tissue of virus-infected animals revealed that the expression of all candidate genes trended towards reduction in male, but not female, *Ube3A* 2xTg brains (Fig. 5B-5G). Among these genes, expression of *lgfbpl1, Rps6ka5* and *Draxin* was significantly decreased in Tg males (Fig. 5B, 5D, 5E). *Pde1c*, *Grik2* and *Pxdn* exhibited a trend of reduced expression, although the changes were not statistically significant (Fig. 5F, 5G). More importantly, ICV injection of LV-GFP-AR in Tg males effectively rescued the expression of these candidate genes to a level comparable to that of WT males (Fig. 5B-5G).

Together, these results showed that AR restoration rescued the expression of ASD-associated ARGs, suggesting that the altered transcriptional activities of AR in the brain may be the key molecular event leading to the male-biased behavioral deficits.

### AR reduction is responsible for male-biased impairments in social preference, repetitive self-grooming and memory

Previous research has established a link between dysregulation of AR and changes in multiple autistic features, such as social behaviors ^51, 62^, learning and memory ^63, 64^, and hyperactivity symptoms ^52, 65^. Therefore, it is intriguing to hypothesize that AR plays a crucial role in mediating the sexually dimorphic autism-relevant behaviors observed in *Ube3A* 2xTg mice. We first sought to determine whether AR restoration by ICV injection of LV-GFP-AR at postnatal day 0 can rescue the impaired social behaviors in *Ube3A* 2xTg mice. In the three-chamber sociability test, when testing social preference by examining interaction time with a stranger mouse versus an empty cage, impaired preference was found only in Tg males not Tg females (Fig. 1C). Similarly, Tg males with LV-GFP injection lost the preference to the stranger mouse compared to WT males with LV-GFP, whereas Tg females with LV-GFP showed normal social preference as compared to WT females with LV-GFP (Fig. 6A, 6B). Surprisingly, ICV injection of LV-GFP-AR rescued social preference in male *Ube3A* 2xTg mice, but had minimal effect on the social preference of female *Ube3A* 2xTg animals (Fig. 6A, 6B). For social novelty, where the test mouse was examined on its preference between the novel and the familiar mouse, LV-GFP-AR failed to rescue the abnormal preference observed in male and female Tg animals (Fig. 6C, 6D). These results indicate that AR reduction plays an essential role in the male-biased deficit in social preference, but does not contribute to deficits in social novelty, which is similarly affected in both male and female Tg animals.

**Fig. 6:**
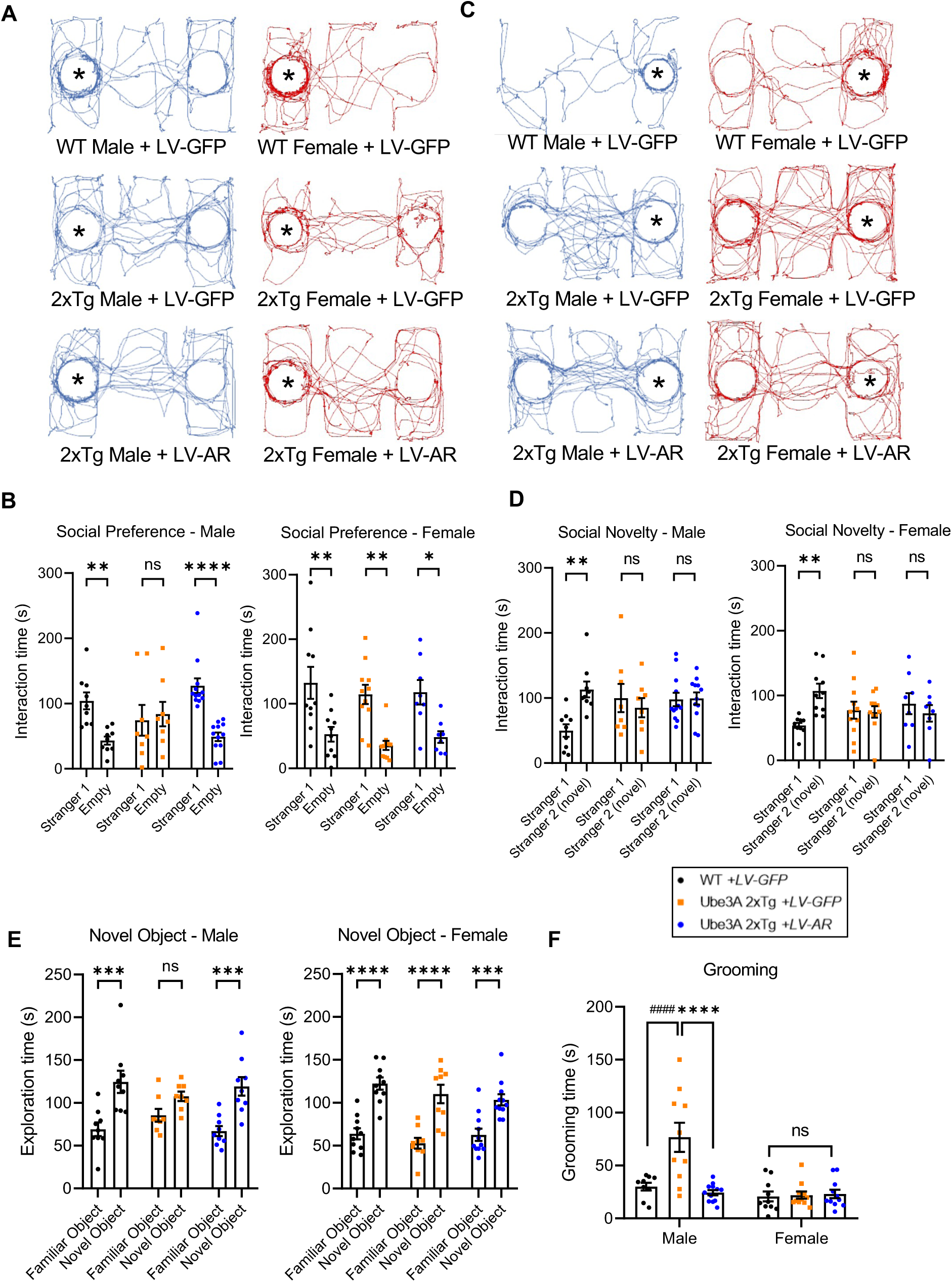
AR restoration in *Ube3A* 2xTg mouse brains rescues social preference, memory, and repetitive self-grooming behaviors in males, without affecting females. **(A-D)** Social behaviors in mice following ICV injection of LV-GFP-AR at postnatal day 0. Three chamber tests were performed at postnatal day 30 to 50. (A) Traces of track paths for social preference test. (B) LV-GFP-AR infection in Tg male rescued its preference for stranger mouse. Tg female showed normal social preference which was not affected by LV-GFP-AR infection. (C) Traces of track paths for social novelty test. (D) LV-GFP-AR infection failed to rescue impaired social novelty in Tg male and Tg female. WT male+LV-GFP: n=9, Tg male+LV-GFP: n=8, Tg male+LV-GFP-AR: n=12, WT female+LV-GFP: n=10, Tg female+LV-GFP: n=11, Tg female+LV-GFP-AR: n=8. **(E)** Long term memory in mice following ICV injection of LV-GFP-AR at postnatal day 0, detected by novel object recognition test at postnatal day 30 to 50. Tg males with LV-GFP-AR displayed better discrimination between the familiar vs. novel object compared to the LV-GFP infected Tg males, indicating successful rescue of memory. Tg female showed normal memory which was not affected by LV-GFP-AR. WT male+LV-GFP: n=9, Tg male+LV-GFP: n=8, Tg male+LV-GFP-AR: n=9, WT female+LV-GFP: n=10, Tg female+LV-GFP: n=9, Tg female+LV-GFP-AR: n=11. **(F)** Repetitive self-grooming behavior was impaired only in Tg males but not females. ICV injection of LV-GFP-AR successfully rescued grooming behaviors in *Ube3A* 2xTg males with no significant effect on *Ube3A* 2xTg females. WT male+LV-GFP: n=9, Tg male+LV-GFP: n=10, Tg male+LV-GFP-AR: n=12, WT female+LV-GFP: n=10, Tg female+LV-GFP: n=11, Tg female+LV-GFP-AR: n=11. Mean ± SEM. *p<0.05; **p<0.01; ***p<0.001; ****p<0.0001. ns, not significant. Two-way ANOVA with Bonferroni’s multiple comparisons test.

We then examined whether AR restoration by LV-GFP-AR could rescue long-term memory by assessing novel object recognition, which was impaired mainly in male *Ube3A* 2xTg mice (Fig. 2C). Quantification of exploration time displayed successful rescue of memory by LV-GFP-AR in *Ube3A* 2xTg males, indicated by a significant increase in the time spent exploring the novel object (Fig. 6E). Tg female mice showed normal memory and were not affected by LV-GFP-AR injection (Fig. 6E). Further, repetitive self-grooming behavior was also differentially affected in *Ube3A* 2xTg animals, with only male Tg mice showing increased grooming (Fig. 1E). Similarly, LV-GFP-AR successfully reversed the grooming behavior back to WT levels, and did not alter the grooming time in Tg females (Fig. 6F).

Together, these results demonstrate that AR restoration in the brain at P0 was able to rescue several male-biased behavioral deficits in Tg animals, without affecting normal behaviors in female Tg mice, indicating that reduced AR signaling in the brain contributes to the expression of autism-relevant behaviors in male mice.

## Discussion

As one of the most common genetic factors in ASD etiology, UBE3A overdosage or hyperactivity correlates with ASD in humans and autistic-like behaviors in mouse models ^14, 16, 17, 19, 23, 24^. By studying the differential effects of UBE3A overexpression in male and female mice, we found that male-predominant behavioral impairments, including deficits in social preference, memory and repetitive grooming behaviors, are associated with the downregulation of androgen receptors. Our findings demonstrate the existence of sexually dimorphic behavioral abnormalities and provide new insight into the molecular underpinnings of pathogenesis in UBE3A-dependent ASD.

Behaviorally, male *Ube3A* 2xTg mice showed deficits in both social preference and social novelty, and were more severely affected in their repetitive self-grooming behaviors, long-term memory, and pain sensitivity as compared to Tg females. Conversely, female *Ube3A* 2xTg mice displayed impairments in social novelty but not social preference, and were more severely impacted in their olfactory sensitivity. Interestingly, both male and female Tg mouse pups showed alterations in communication as indicated by their mother-seeking USVs induced by maternal separation. While the male Tg mice tend to call more, female Tg mice tend to use more complex syllables. These behaviors are consistent with the syndromes observed in patients with excessive UBE3A activity, including typical autism-relevant behaviors such as social and communication deficits, intellectual disability, as well as sensory disorders ^16^.

Recent studies have proposed diverse mechanisms underlying the sexually dimorphic expression of ASD. Sex-differential factors such as sex chromosomes and sex hormone signaling may function as sex-specific risk factors or protective factors ^7, 70^. For instance, fetal testosterone during development has been proposed as one potential factor that may contribute to the male-biased features, like impaired social behaviors and restricted interests ^71, 72^. Recent research has also identified non-sex-differential factors, such as transcriptomic alterations. The downregulation of neuronal genes and the upregulation of glial genes in males with autism may contribute to the sex imbalance in ASD ^73^. Some studies suggest that females require a higher mutation load to exhibit the diagnostic level of autism-relevant phenotypes, leading to the development of the “female protective model” ^8, 74, 75^. Furthermore, sexually dimorphic mechanisms related to ASD may involve not only transcriptomic changes, but also other mechanisms, such as synaptic transmission and neuronal activity ^76–78^. Despite these findings, understanding of the biological mechanisms remains limited.

By compiling case reports, we find that approximately 70% of dup 15q11-13 ASD cases are males ^79–85^. The male-predominant behavioral deficits in *Ube3A* 2xTg mice are consistent with this higher incidence of ASDs reported in males with the 15q11-13 duplication than in females. Previous studies on *Ube3A* 2xTg mice often limit their data analysis to comparing the differences between WT and Tg, but not male vs. female animals ^23, 24^. Interestingly, upon combination of the male and female data with a near 1:1 ratio, our behavioral data are in line with the findings from previous research ^23, 24^. Specifically, *Ube3A* 2xTg animals exhibit notable alterations in social preference, social novelty, vocalization, and repetitive self-grooming behaviors (Fig. S2). Of note, memory functions and pain sensitivity appear to be largely intact when the data from both sexes are combined (Fig. S3D, S3F-S3I), indicating that deficits in memory and pain sensitivity observed in male Tg mice (Fig. 2C, 2G, 2K) can become indiscernible when that data is diluted by the normality in female Tg mice (Fig. 2C, 2G, 2K). Similarly, when male and female results were combined, olfactory sensitivity, which was mainly impaired in Tg females (Fig. 2N), did not show a significant difference between WT and *Ube3A* 2xTg mice (Fig. S3E). Obviously, sex-orientated data analysis is important to reveal subtle phenotypes hidden in a general population of both sexes.

The crucial role of AR in brain development and function has been well documented. AR is strongly expressed in the brain, and in fact, male and female mouse brains have comparable levels of ARs at most time points during development, except postnatal day 7 ^86^. However, the ligands of ARs, androgens, differ significantly between male and female brains ^87^. Both testosterone and 5*α*-dihydrotestosterone are markedly more abundant in male brains than in female brains, resulting in much higher activities of AR signaling in male brains ^87^. In line with this, we find that a similar reduction of AR caused more severe disturbances in male compared to female mice in gene regulation and behaviors.

The androgen system regulates a range of behaviors, including male sexual behaviors, social behaviors, as well as learning and memory ^51, 62, 64, 88, 89^. In addition, ARs have been shown to increase the neural progenitor cell pool in the developing cortex, further indicating its vital role in neurodevelopment ^48^. Therefore, dysfunction of ARs is expected to interfere with brain development. In fact, several studies have linked AR dysregulation to ASD. Specifically, *AR* itself and at least seven AR target genes have been identified as ASD genes ^5^. Altered AR activity has also been implicated in the hyperactivity symptoms of ASD, such as ADHD ^52, 65^, which is a common comorbidity found in 30% of autism patients ^52^. Our findings further confirmed the crucial role of AR in regulating autism-relevant behaviors, including social preference, repetitive self-grooming and memory. Continued investigation into the role of ARs in neurodevelopment and neurological diseases may lead to the development of novel therapeutics, such as the potential of using AR agonists as a treatment strategy for UBE3A-dependent ASD.

### Limitations of the study

Intracerebroventricular injection of LV-GFP-AR virus was conducted in P0 mice, which limits the ability to assess the effects of AR replenishment on prenatal brain development and its potential influence on sex differences and autistic behaviors. Future studies involving prenatal replenishment of AR expression will help determine the roles AR reduction may play during embryonic stages.

## Materials and Methods

### Antibodies

The following antibodies were used in the indicated dilution or amount: Mouse anti-AR (WB: 1:500; IF: 1:200; IP: 1.5 µg; Santa Cruz Biotechnology; Cat# sc-7305). Mouse anti-ERα (WB: 1:500; Invitrogen; Cat# MA1-80216). Rabbit anti-ERβ (WB: 1:500; Invitrogen; Cat# PA1-311). Mouse anti-UBE3A (E6AP; WB: 1:1000; IF: 1:200; Sigma-Aldrich; Cat# E8655). Mouse anti-Ubiquitin (WB: 1:500; Invitrogen; Cat# 14-6078-82). Mouse anti-GAPDH (WB: 1:5000; EMD Millipore; Cat# MAB374). Mouse anti-α tubulin (WB: 1:5000; Sigma-Aldrich; Cat# T9026).

### Animal care and use

Ube3A conventional BAC overexpressing mouse models on the FVB background (*Ube3A*-Tg 2x, strain #019730) ^23, 24^ and FVB/NJ WT mice (strain #001800) were used in this study. All animal procedures were conducted in compliance with the policies of the Institutional Animal Care and Use Committee (IACUC) at Boston University. *Ube3A* 2x transgenic (2xTg) animals were obtained by mating heterozygous males and heterozygous females, expected to produce a 1:2:1 (WT: 1xTg: 2xTg) offspring ratio. WT and homozygous *Ube3A* 2xTg mice from the litters were used for the experiments, whereas the 1xTg mice were kept for further breeding. Therefore, the number of independent litters for the WT group and that for the 2xTg group used in the experiments is similar. Both male and female mice were used for behavioral tests and tissue collection. Most mice underwent multiple behavioral tests, with individual tests being separated by at least three days to prevent interference. Animals naïve to behavioral tests were used for the majority of the brain work, except those used for confirmation of virus expression and the rescue effect on ARGs.

### Behavioral tests

For all behavioral tests, male and female mice were trained and tested in parallel. During all behavioral tests, except the Barnes maze spatial memory test, the testing room was dimly lit by a small desk lamp in the corner to enable the experimenter to maneuver. For the Barnes maze spatial memory test, bright ceiling lamps were used as an aversive stimulus.

### Three-chamber social test

The three-chamber social test was performed as described in our previous study ^90^. A three-chambered box was made from plastic board that is 0.75 inches thick and measures 65 x 28 x 28 cm. The walls to the center chamber had cut-out doors that are 10 x 10 cm in size allowing movement between chambers. The two side chambers contained small wire cages to house the social stranger and/or novel mice. Before the test, mice were habituated to the apparatus with empty cages in each side chamber for 3 days, with each session lasting 5 min, allowing them to move freely within the apparatus. Social Preference Test: mice were singly placed into the middle chamber, with the doors blocked by plastic boards and a stranger mouse was placed into one of the side chambers under the wire cage. The doors were unblocked, and the test mouse was allowed to move freely within the apparatus for 5 min. Social Novelty Test: the test mouse was then returned to the middle chamber, the doors were blocked again. A second mouse (novel mouse) was introduced into the other side chamber. The center doors were unblocked and the test mouse was allowed to move freely within the apparatus for another 5 min. The entire apparatus was wiped with 70% ethanol between mice to eliminate odor cues between animals. Video recordings were captured with a Logitech c920 webcam during the test and the time spent interacting with each mouse or empty cage (nose ≤ 2 cm) and locomotion tracks were scored using TrackMo, an open-source animal tracking toolbox available at https://github.com/zudi-lin/tracking_toolbox.git, with the analyzer blinded to the genotype of the animals.

### Ultrasonic vocalization (USV) recording

Ultrasonic vocalization recording of pup isolation calls was performed as described in our previous work ^90^. On the test day, pups were separated from the parents and littermates, and isolated in a sound-proof cylindrical plastic container. Mouse vocalizations were recorded on P5, P7 and P9, in a random order for each litter, using a CM16/CMPA microphone (Avisoft Bioacoustics) positioned 15 cm above the pups. The recording chamber was cleaned with 70% ethanol and dried between each pup. USV signals were recorded for 5 min at a sampling rate of 300 kHz in 16-bit format. The microphone was connected to a preamplifier UltraSoundGate 116Hb (Avisoft Bioacoustics) and digitized sonograms were stored in a computer. Recordings were analyzed using SASLab Pro 5.2.12 (Avisoft Bioacoustics). Spectrograms were generated using fast Fourier transform (256 FFT length, 100% frame, FlatTop window, and 50% window overlap) and a high-pass filter was applied to eliminate background noise <25 kHz. USVs were detected automatically (threshold: −47 ± 10 dB, hold time: 7 ms) followed by manual inspection to ensure accuracy. Several acoustic parameters of calls were measured including number of calls, mean call duration, total time spent calling, peak frequency, and peak amplitude. Call syllables were classified based on their acoustic features: simple (short, flat, upward, downward, chevron, U shape) and complex (modulated, frequency jump, multiple jumps, harmonic).

### Self-grooming behavior test

The self-grooming behavior test was performed as described in our previous work ^91^. Each test mouse was placed individually into a regular housing cage with a fresh thin layer (∼0.5 cm) of wood chip beddings and was allowed to acclimate for 20 mins. The mice were then video recorded for 10 mins from the side view of the cage by a camera. The total time spent on grooming was measured.

### Barnes maze spatial memory test

The Barnes maze spatial memory test was performed as described in our previous work ^90^. The Barnes maze is a circular maze constructed from a plastic board with a diameter of 48 inches and a thickness of 0.75 inches. Twenty holes with a diameter of 2 inches were evenly spaced around the perimeter of the maze, each with a distance of 1 inch to the edge. The maze was mounted on a 30-inch-high pedestal and could rotate at its center. The escape cage was a black plastic box with a ramp connected beneath the escape hole for easy access. During testing, four bright ceiling lamps and an alarm were used as aversive stimuli. To prevent olfactory cues, the maze and escape cage were cleaned with 70% ethanol between each testing session, and the maze was randomly rotated after each mouse. During the test, the mice were habituated to the maze on day 1. For each session, one mouse was placed in the center of the maze, covered with an opaque cardboard chamber for 15 s, and then slowly guided to the escape hole with a glass beaker. Each mouse was given 3 min to enter the escape hole on its own, but if they failed to do so, they were nudged gently into the hole. Afterward, the mouse was allowed to stay in the escape hole for 2 min. The ceiling lights and white noise remained on while the mouse was exploring the maze and turned off immediately after they entered the escape hole. This procedure was repeated for all mice. On days 2–5, the mice were trained to ensure a strong memory for the escape hole (4 times/mouse/day for 4 days, 16 times in total) and to learn to enter the escape hole by themselves. During each training trial, the mice were allowed to stay in the escape cage for 1 min. Memory of the escape hole location was probed 24 hours and 5 days later. During the memory retention tests, the escape hole was sealed and spatial memory for the subject mice was tested. Video recordings were captured for 4min. The number of errors committed prior to locating the target hole (primary errors) and traces of locomotion were scored by TrackMo.

### Novel object recognition (NOR) test

The novel object recognition test was performed as described in our previous work ^92^. The NOR protocol comprises three sessions: habituation, training, and test sessions. During habituation, each mouse exposed to an open-field apparatus twice a day for 3 consecutive days, with each session lasting 10 min. On day 4, two identical objects (falcon tissue culture flask filled with beddings) were placed in two corners of the arena and mice were allowed to explore freely for 10 min. 4 hours or 24 hours later, during the test sessions, one object was replaced by a novel object (tower of Lego bricks). Video recordings were captured for 5 min, and exploration time for each object and traces of locomotion were scored by TrackMo.

### Hot plate test

To assess the pain sensitivity of mice, the hot plate test was conducted following the protocol detailed in our previous study ^92^. Prior to the test, mice were habituated to the non-heated hot plate surface and a covered 5L glass beaker for 2 days (15 min/day). During day 3, the hot plate temperature was set to 55°C. Mice were placed on the hot plate and covered with a 5L glass beaker. The latency to response was recorded when the hind paw lick occurred.

### Olfactory detection threshold test

The olfactory detection threshold test was performed as described in previous protocols with minor modifications ^93^. On day 1, two filter papers were placed at two corners of a regular housing cage, one with a vanilla solution at the lowest concentration and the other with a filter paper soaked in water (vehicle). The test mouse was allowed to freely explore for 3 min, and the time spent on each filter paper was recorded. During day 2-4, the session was repeated once a day, but using vanilla solution with a 100 times higher concentration each day. If the mouse cannot detect the odorant, it was expected to spend an equal amount of time sniffing the vanilla paper and the vehicle. The percentage of odorant sniffing was calculated using the following equation: the percentage of odorant sniffing = time spent on odorant / (time spent on odorant + time spent on vehicle) X 100.

### Open-field test

The open-field test was performed as described in our previous work ^90^. The open-field test was conducted in a chamber that measured 28 x 28 x 28 cm and was made of 0.75-inch-thick plastic board. The chamber was placed on a large black plastic board, and the lights in the testing room were turned off except for a small desk lamp in the corner allowing the experimenter to see. Mice were habituated to the testing room for 5 min/day for 3 days. On the test day, each mouse was placed individually in the center of the box and allowed to freely explore the chamber. Video recordings were captured during the test and the time spent in the center, velocity and track lengths were quantified blinded to the genotype of the animals.

### Primary neuron culture

Primary cortical neurons were prepared from embryonic day 18 rat fetuses as described previously ^90^. Timed pregnant Sprague–Dawley rats were purchased from Charles River Laboratories Inc. Cortical brain tissues were dissected and digested with papain (0.5 mg/mL in Hanks balanced salt solution, Sigma-Aldrich; cat. #4762) for 20 min at 37°C, then gently triturated in a trituration buffer containing 0.1% DNase (cat. #PA5-22017 RRID: AB_11153259), 1% ovomucoid (Sigma-Aldrich; cat. #T2011), 1% bovine serum albumin (Sigma-Aldrich; cat. #05470) in DMEM to fully dissociate neurons. Dissociated neurons were counted and plated onto 18mm circular coverslips (Carolina Biology Supply, cat. #633013). Coverslips had been coated in poly-l-lysine (Sigma-Aldrich; cat. #P2636; 100 μg/ml in borate buffer) overnight at 37°C, then washed three times with sterile deionized water and left in plating medium [DMEM supplemented with 10% fetal bovine serum (Atlanta Biologicals; cat. #S11550), 5% horse serum (Atlanta Biologicals; cat. #S12150), 31 mg of l-cysteine, 1% penicillin/streptomycin (Corning; cat. #30–002-Cl), and l-glutamine (Corning; cat. #25–005-Cl)] before cell plating. The day after plating, plating medium was replaced by feeding medium [neurobasal medium (Gibco; cat. #21103049) supplemented with 2% Neurocult SM1 Neuronal Supplement (StemCell Technologies; cat. #05711), 1% horse serum (Atlanta Biologicals; cat. #S12150), 1% penicillin/streptomycin (Corning; cat. #30-002-CI), and l-glutamine (Corning; cat. # 25-005-CI)]. One week after plating, *5-fluorodeoxyuridine* (10μM; Sigma-Aldrich; cat. #F0503) was added to the medium to inhibit glial growth. All cells were maintained in a humidified incubator containing 5% CO_2_ at 37°C.

### Immunocytochemistry of cultured neurons

Culture neurons were washed twice in ice-cold PBS and fixed for 10 min in a 4% paraformaldehyde solution at room temperature. Cell membranes were then permeabilized for 8 min in 0.3% Triton X-100 (Fisher Biotec; cat. #BP151-100) in PBS. Cells were rinsed three times in PBS, and blocked in 10% goat serum for 1 h. Primary antibodies (in 5% goat serum PBS) were added, and the cells were incubated overnight at 4℃, washed twice with PBS and incubated with appropriate Alexa Fluor-conjugated fluorescent secondary antibodies (1:400, Life Technologies) for an additional hour. Cells were then rinsed (with Hoechst 1:2000 in the first rinse) and mounted to microscopy glass slides with Prolong Gold anti-fade mounting reagent (Invitrogen; cat. #P36930) and stored at 4℃ in dark for subsequent imaging.

### Neuronal and HEK293T cell transfection

Neurons were transfected with Lipofectamine 2000 (Invitrogen, cat. #11668019) and the target plasmids on DIV4 according to the manufacturer’s instruction. Each coverslip was transfected with 1 μg of either GFP or HA-UBE3A plasmid (Addgene, cat. #8648), and 1 μL Lipofectamine 2000. The plasmid and Lipofectamine 2000 were diluted separately in 50 μL DMEM in 1.5 mL Eppendorf tubes and incubated for 5 min. Then solutions in two tubes were mixed together and incubated at room temperature for another 20 min to form the transfection complex. Thereafter, the transfection complex was gently added into the 12-well plate with 1 coverslip/well in a drop-wise manner. The neurons were incubated with the transfection complex at 37 °C for 3 h before the medium was removed and replaced with feeding medium. On DIV8 (4 d following transfections), neurons were fixed for immunocytochemistry or lysed for biochemical analysis. HEK cell transfections were performed using the polyethylenimine (PEI) transfection reagent (Polysciences; cat. #23966) with a 3:1 PEI-to-DNA ratio when the cells reached 70% confluency. HEK cells were lysed and collected 48 h post-transfection.

### Immunoprecipitation

For ubiquitination immunoprecipitation assays, cultured cortical neurons, HEK cells, or brain tissues were rinsed with cold PBS and lysed on ice in 200 μl of modified radioimmunoprecipitation assay (RIPA) lysis buffer (50 mM Tris-HCl, pH 7.4, 150 mM NaCl, 1% NP-40, 1% sodium deoxycholate) supplemented with cOmplete protease inhibitor (Roche; cat. #11873580001). Cell or brain lysates were further solubilized by sonication and 1 h rotation at 4℃. After rotation, lysates were centrifuged at 12000g, 20 min, 4℃ to collect supernatant. A small portion of each sample was saved as a total cell lysate while the reminder was adjusted to 500μl with more RIPA buffer for immunoprecipitation. Protein A-Agarose beads (Santa Cruz Biotechnology; cat. #sc-2001) were added to the lysates along with antibodies against AR (Santa Cruz Biotechnology; cat. # sc-7305) and samples were incubated overnight for 12–16 h on rotation at 4°C. Agarose beads were washed three times with cold RIPA buffer, resuspended in 2X Laemmli sample buffer, and denatured at 95°C for 10 min before being subjected to Western blotting.

### Western blot

Cultured neurons were lysed in 2X Laemmli sample buffer (4% SDS, 10% 2-mercaptoethanol, 20% glycerol, 0.004% bromophenol blue, 0.125 M Tris HCl) and boiled for 10 min at 95°C before being stored at −20°C for later use. Mouse brain tissues were dissected on ice immediately after sacrificing animals, and then lysed on ice using RIPA lysis buffer. Samples were further solubilized by sonication and 1 h rotation at 4°C. Lysates were then centrifuged at 12000g, 20 min, 4°C to collect supernatants, which were then subjected to a BCA assay according to the manufacturer’s protocol (Pierce, cat. #23225) to determine protein concentrations. Protein levels were normalized using RIPA lysis buffer. Samples were then mixed with 2X Laemmli sample buffer and boiled for 10 min at 95°C to prepare for SDS-PAGE.

Standard SDS-PAGE procedures were used to separate proteins of interest in the cell or brain lysates. Proteins were transferred to PVDF membranes (Bio-Rad, cat. #1620177) and blocked for 1 h in 5% milk in PBS. Membranes were then probed for different targets with the appropriate primary antibodies overnight at 4°C in Tris buffered saline supplemented with 0.05% Tween (TBST). Membranes were washed 3 times in TBST, followed by incubation with the appropriate secondary antibody for 1 h. After a further three washes with TBST, immunoblots were visualized using a chemiluminescence detection system (Sapphire Biomolecular Imager, Azure biosystems, CA, USA) and analyzed using ImageJ.

### Immunohistochemistry of brain slices

As described in our previous work ^90^, before removing the brain from the skull, mice were subjected to transcardial perfusion with ice-cold PBS. Brains were fixed in 4% paraformaldehyde for 4–6 h and then cryoprotected in 30% sucrose/PBS until sinking to the bottom. To prepare for sectioning, the dehydrated brains were frozen in optimal cutting temperature-embedded medium (Tissue-Tek, cat. #25608-930) and sliced into 20 μm sections using a CM1850 cyrostat (LEICA Biosystems) at −20°C. Sections were immediately mounted onto SuperFrost microscope slides (Fisher Scientific, cat.# 12-550-15) and stored at −80°C for future use. Prior to immunostaining, sections were rinsed with PBS for at least 2 h. Sections were then blocked in 5% goat serum, 0.3% Triton X-100/PBS for 1 h at room temperature, followed by incubation with properly diluted primary antibodies overnight at 4°C. The following day, sections were washed three times with PBS and incubated with the appropriate Alexa Fluor-conjugated secondary antibodies (1:250, Life Technologies). Brain slices were then washed three times with PBS, with the first wash containing Hoescht (1:2000, Thermo Fisher Scientific, cat. # 62249). Sections were then mounted with ProLong-Gold mounting medium (Invitrogen, cat. #P36930), left to dry at room temperature overnight, and stored at −20°C in dark before imaging.

### Viral constructs preparation and virus infection

GFP-fused AR was cut from pEGFP-C1-AR plasmid (Addgene, cat. #28235) and inserted into the pFUW lentiviral (LV) vector, to generate pFUW-GFP-AR construct. To package LV particles, HEK293T cells at 70% confluency in 15-cm culture plates were transfected with pFUW-GFP-AR, along with viral packaging and envelope proteins (pRSV/REV, pMDLg/RRE, and pVSV-G) using polyethylenimine transfection reagent (Polysciences; cat. #23966). 48 h post-transfection, cell medium was filtered through a 0.45 μm filter, and PEG-it virus precipitation solution (1:5, System Biosciences; cat. # LV810A-1) was added and kept at 4°C overnight. On the next day, the solution was centrifuged (1500 rpm, 30 min, 4°C), and the pellet was resuspended with 100 μl cold PBS. Aliquots of the virus were stored at −80°C until use. For in vitro assays, neurons were infected with lentivirus at DIV6, and the medium was replaced with feeding medium 2 d later.

### Intracerebroventricular brain injection of virus in neonatal mice

Bilateral intracerebroventricular (ICV) injections were performed in mouse pups on postnatal day 0. Newborn mice were cryoanesthetized and placed on a cold metal plate. A 30-gauge needle (Hamilton, cat. # 7803-04) and 10 μL Hamilton syringe (Hamilton, cat. # 7653-01) were used for injection. To pierce the skull, a separate needle was employed at a site that is roughly two-fifths of the distance between the lamda and the eye. Each cerebral ventricle was slowly injected with 2 μL of either pFUW-GFP or pFUW-GFP-AR virus mixed with 1% Fast Green Dye. Following injection, neonatal mice were kept with their parents until they were weaned. On postnatal day 30, all mice were subject to behavioral assessments before being euthanized for biochemical and histological analysis.

### Real-time PCR

Total RNA from prefrontal cortical area was extracted using the RNeasy® Mini Kit (Qiagen, cat. # 74104) following the manufacturer’s instructions. cDNA was reverse transcribed from 500 ng of total RNA for each sample using HiFiScript cDNA Synthesis Kit (CoWin Biosciences, cat. # CW2569M) according to the manufacturer’s protocol. cDNA samples were then used as a template for further mRNA level analysis.

For qPCR, 1 μL of the reverse transcribed cDNA was used as a template and mixed with primer pairs:

AR (5′- GATGGTATTTGCCATGGGTTG-3′, 5′- GGCTGTACATCCGAGACTTGTG-3′);

lgfbpl1 ((5′-CTGACCATGAGACCACATCCTG-3′, 5′-ACTGAGCCTCTCCAATGGCGTT-3′);

Pde1c (5′-CAGTCATCCTGCGAAAGCATGG-3′, 5′-CCACTTGTGACTGAGCAACCATG-3′);

Rps6ka5 (5′-TGGTCCATAGCACCTCTCAGCT-3′, 5′-CTCTCCGCCATTCAGAAGTTCC-3′);

Draxin (5′-GTGGCAGAGAACACAAGAGACG-3′, 5′-GGTCTTCAGAGGGTTCCACCTT-3′);

Grik2 (5′-CTGACTCAGGTTTGCTGGATGG-3′, 5′- CAAGGAGCTGACTGTCATCTGG-3′);

Pxdn (5′-GTTCAGCATGGCTTGATGGTGG-3′, 5′-AGCCTGACAGGTTGGCGATGAG-3′);

10 μL reactions were prepared with UltraSYBR Mixture kit (CoWin Biosciences, cat. #CW2602F) according to the manufacturer’s protocol and placed in Applied Biosystems 7900HT Fast Real-Time PCR system for real-time monitoring.

### RNA Sequencing

Brain tissues from three animals in each group (WT male, WT female, Tg male, Tg female) were dissected and preserved in RNAlater™ Stabilization Solution (Invitrogen, cat. #AM7020) before being sent to commercial RNA-sequencing services provided by BGI genomics. RNA-seq datasets were generated and analyzed by BGI genomics to identify differentially expressed genes and gene ontology enrichment. Datasets from *Ube3A* 2xTg males were compared with those from WT males, and datasets from *Ube3A* 2xTg females were juxtaposed with those from WT females.

### Statistics and reproducibility

Graphics and statistical analysis were performed using GraphPad Prism 8.0 (GraphPad Software). Data of genotypes (for combined results of both males and females, or in vitro assays) were analyzed using an unpaired two-tailed Student’s t-test, with genotype as a factor. Comparisons involving two independent variables, such as sexually dimorphic changes between WT and *Ube3A* 2xTg animals, were analyzed by two-way analysis of variance (ANOVA). Three-way ANOVAs were used for comparisons involving three independent variables, each with at least two levels: interaction items (stranger mouse vs. empty cage, stranger mouse vs. novel mouse, or familiar object vs. novel object), sex (male vs. female), and genotype (WT vs. Tg). Post hoc multiple comparisons were corrected using Bonferroni’s method. Animals that did not exhibit normal behavior or health at the onset of the experiment, or showed signs of injury and sickness during experiments, were excluded. Following statistical analysis, individual data points with a value larger or smaller than 2 standard deviations were considered outliers and removed from the data pool. All results are presented as mean ± s.e.m. (standard error of mean), and *p < 0.05 was considered statistically significant. All statistics and sample sizes are reported in Supplementary Table 1.

## Data availability

Data used in the analysis of this study are available from the corresponding author upon reasonable request.

## Acknowledgements

We would like to thank the Man Lab members for helpful discussion. We thank KathrynAnn Odamah and Dr. Mike Baum for their comments on the manuscript. This work was supported by NIH grants R01 MH079407, R01 MH130600, R21 MH133014 and the Harvard/MIT Joint Research Grants Program in Basic Neuroscience (HYM).

## Author Contributions

HYM and YT contributed to the conception of the study; YT and HQ performed the behavioral experiments; YT performed the biochemical assays and rescue experiments; LQZ contributed to the RNA-sequencing analysis; YT performed the data analysis; HYM guided the experiments and assisted the techniques; HYM and YT wrote the manuscript.

## Declaration of Interests

The authors declare no competing interests.

---

**Fig. S1:**
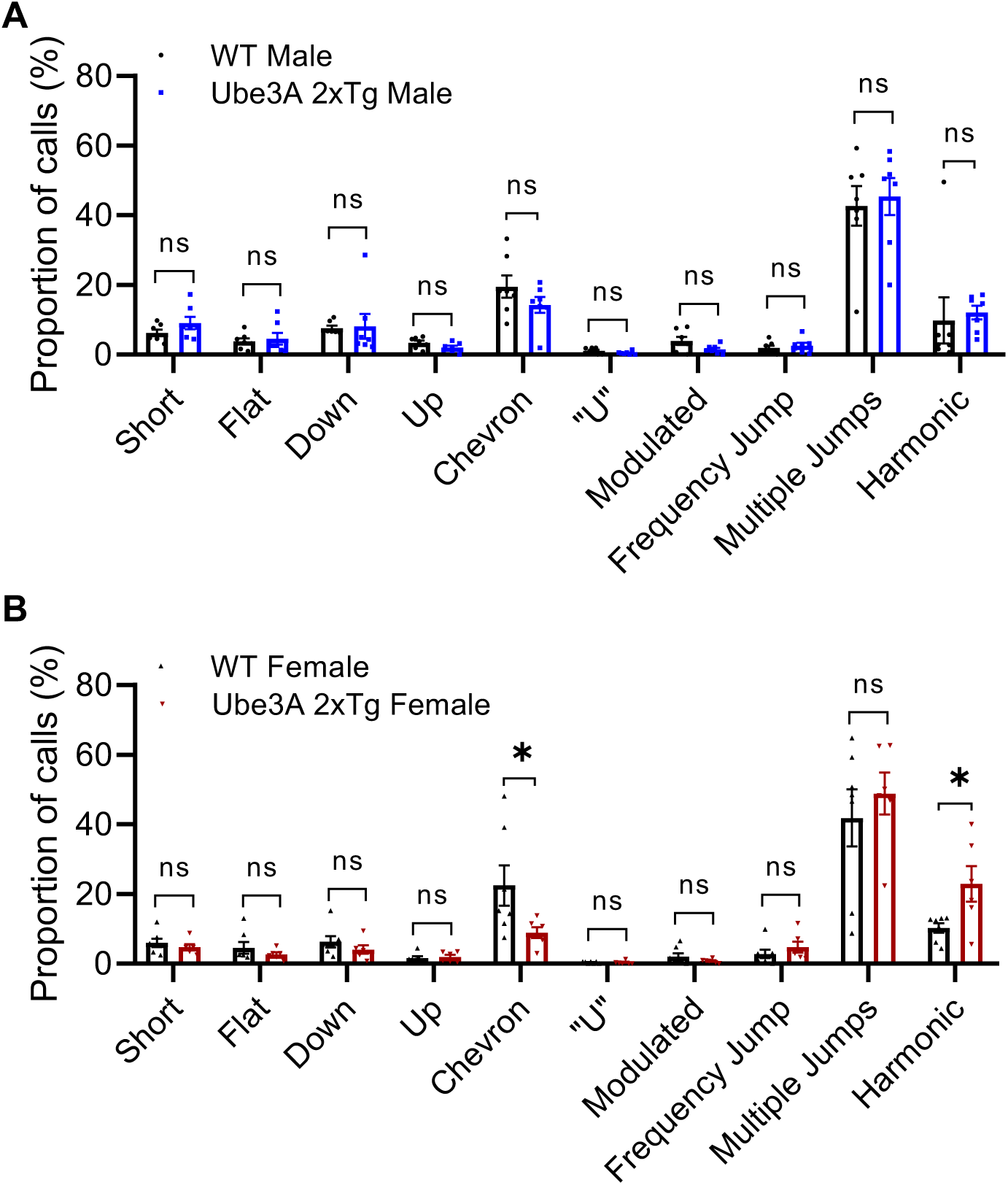
Sexually dimorphic changes in the call syllables of USV in male and female *Ube3A* 2xTg mice. **(A)** Call syllable classification of USV showed no change in *Ube3A* 2xTg males compared to WT males at P5. **(B)** *Ube3A* 2xTg female mice made significantly fewer chevron calls and more harmonic calls compared with WT female mice at P5. n=7 WT male; n=7 Tg male; n=7 WT female; n=6 Tg female. Mean ± SEM. *p<0.05. ns, not significant. Two-way ANOVA with Bonferroni’s multiple comparisons test.

**Fig. S2:**
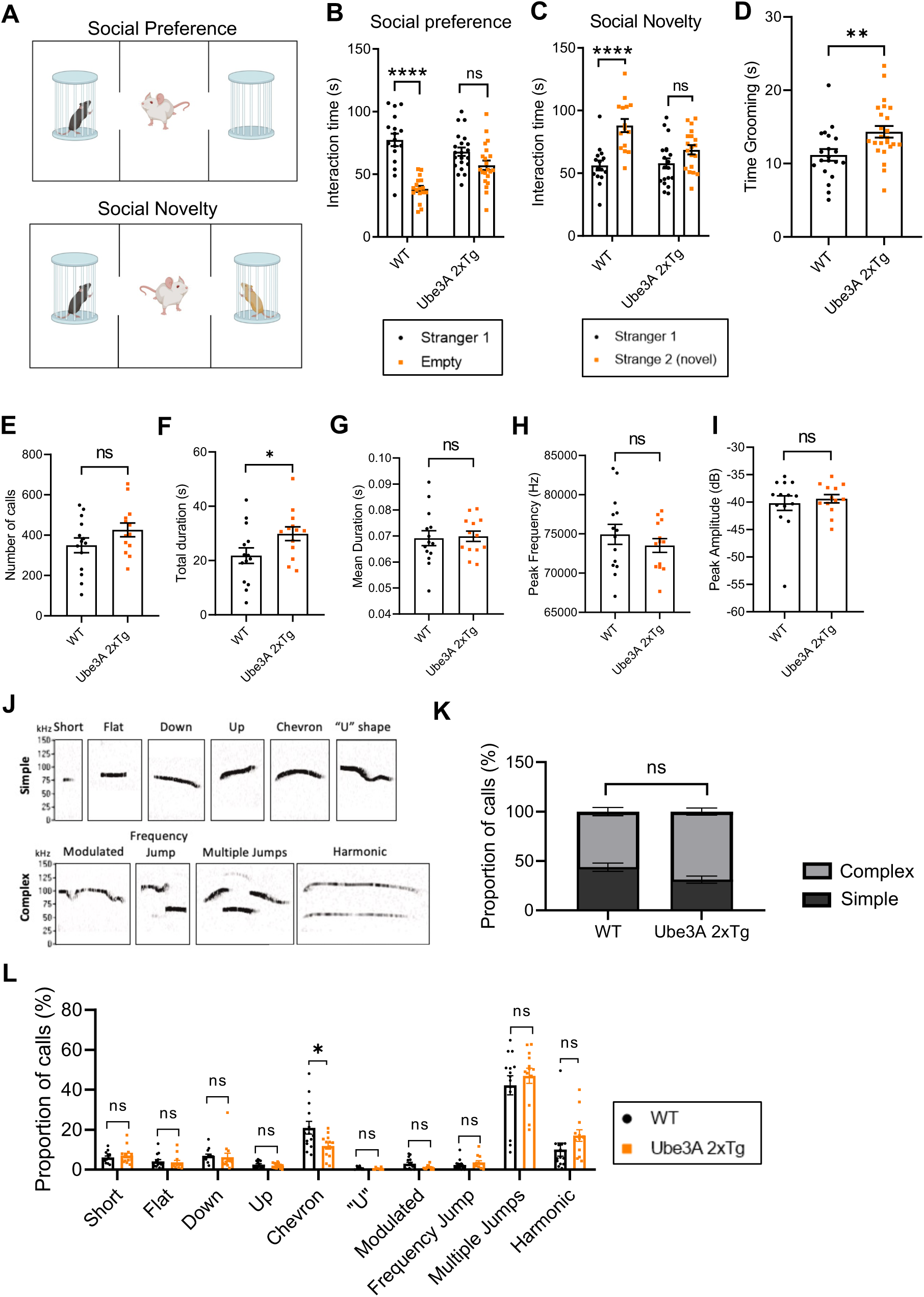
Combining male and female results, *Ube3A* 2xTg mice display impairments in social behaviors, repetitive self-grooming behaviors, and ultrasonic vocalizations. **(A)** The paradigm for the three-chamber social test. For social preference (top), an unfamiliar mouse was placed into either of the side chambers and the test mouse was allowed to move freely in the apparatus. For social novelty (bottom), a novel mouse was placed into the remaining empty chamber, and the test mouse was allowed to interact with both mice. **(B,C)** Quantification of the interaction time showed a decrease in preference for the stranger mouse (B) and the novel mouse (C) in *Ube3A* 2xTg mice. **(D)** *Ube3A* 2xTg mice showed increased grooming behavior. **(E-I)** Quantifications of the number of calls (E), total call duration (F), mean call syllable duration (G), peak frequency (H), and peak amplitude (I) for USVs recorded on postnatal day 5. *Ube3A* 2xTg mice displayed longer total call duration than WT mice. **(J)** Representative calls of each type used in syllable characterization. **(K,L)** WT and *Ube3A* 2xTg animals displayed similar usage of syllabus at P5, except that *Ube3A* 2xTg animals made fewer chevron calls. **In (B)**, n=17 WT, n=21 *Ube3A* 2xTg. **In (C)**, n=15 WT, n=19 *Ube3A* 2xTg. **In (D)**, n=20 WT, n=24 *Ube3A* 2xTg. **In (E-L)**, n=14 WT, n=13 *Ube3A* 2xTg. Mean ± SEM. *p<0.05; **p<0.01; ****p<0.0001. ns, not significant. In (B), (C), (K), (L) Two-way ANOVA with Bonferroni’s multiple comparisons test; in (D), (E-I) Unpaired two-tailed t test.

**Fig. S3:**
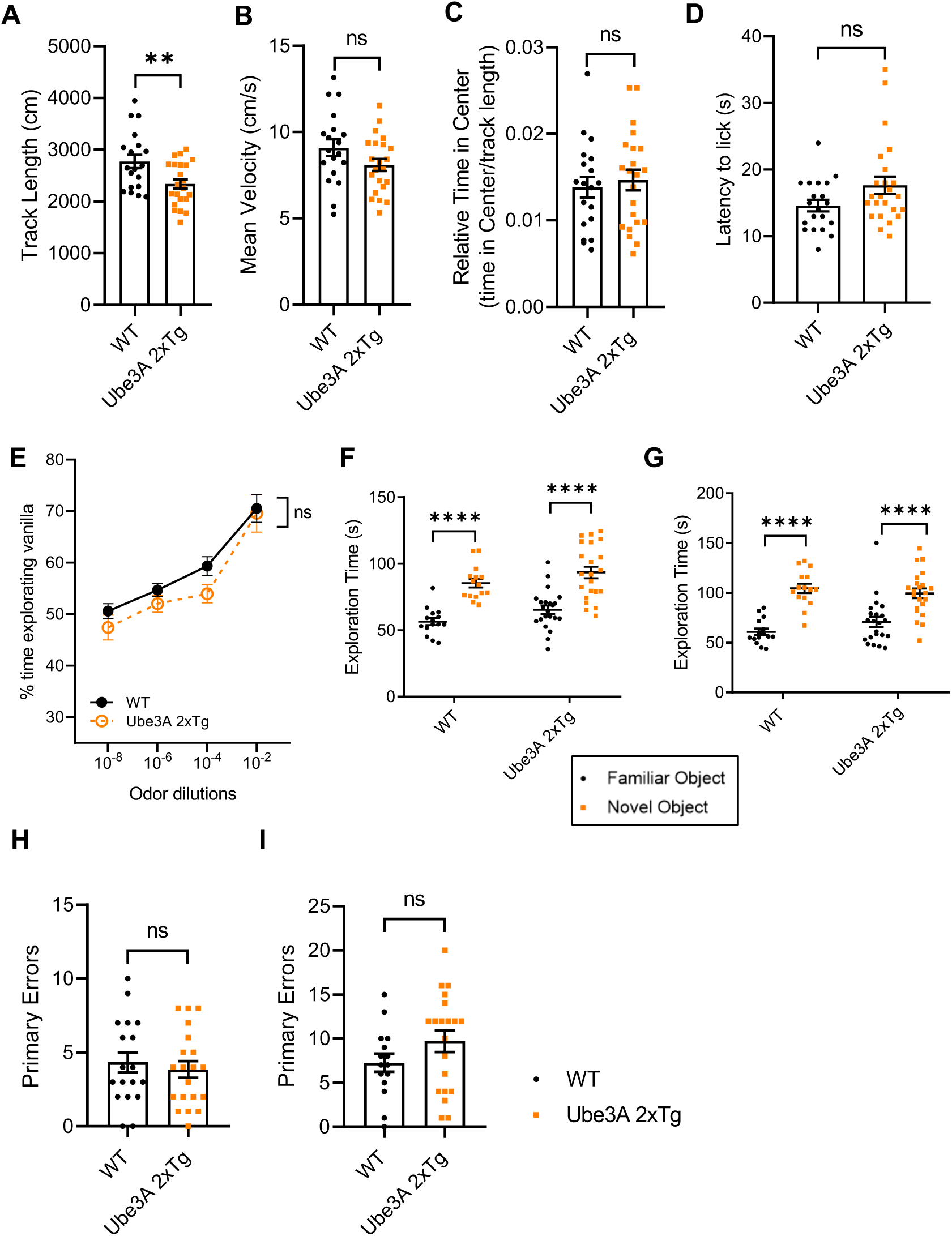
Combining male and female results, *Ube3A* 2xTg mice display reduced activity, normal pain and olfactory sensitivities, and normal memory. **(A-C)** During open field test, *Ube3A* 2xTg animals showed decreased track lengths (A), normal mean velocities (B) and no change in relative time of stay at the center (C). n=19 WT, n=22 *Ube3A* 2xTg. **(D)** Test mice were placed on 55℃ hot plate and their latency to lick hind paws was recorded. *Ube3A* 2xTg mice displayed normal pain sensitivity compared to WT. n=20 WT, n=24 *Ube3A* 2xTg. **(E)** Test mice were exposed to vanilla at varied concentrations. Time spent exploring vanilla vs. water was recorded. The olfactory sensitivity of 2xTg mice was similar to that of WT mice. n=19 WT, n=20 *Ube3A* 2xTg. **(F,G)** During novel object recognition test, *Ube3A* 2xTg animals showed normal exploration time of familiar object vs. novel object 4 h (F) and 24 h (G) post training. n=15 WT, n=22 *Ube3A* 2xTg. **(H,I)** During Barnes spatial memory maze test, primary errors made to find the escape hole 24 h (H) and 5 d (I) after the last training were similar between WT and *Ube3A* 2xTg animals. In (H), n=18 WT, n=20 *Ube3A* 2xTg. In (I), n=15 WT, n=20 *Ube3A* 2xTg. Mean ± SEM. **p<0.01; ****p<0.0001. ns, not significant. In (E-G) Two-way ANOVA with Bonferroni’s multiple comparisons test; in (A-D), (H-I) Unpaired two-tailed t test.

**Fig. S4:**
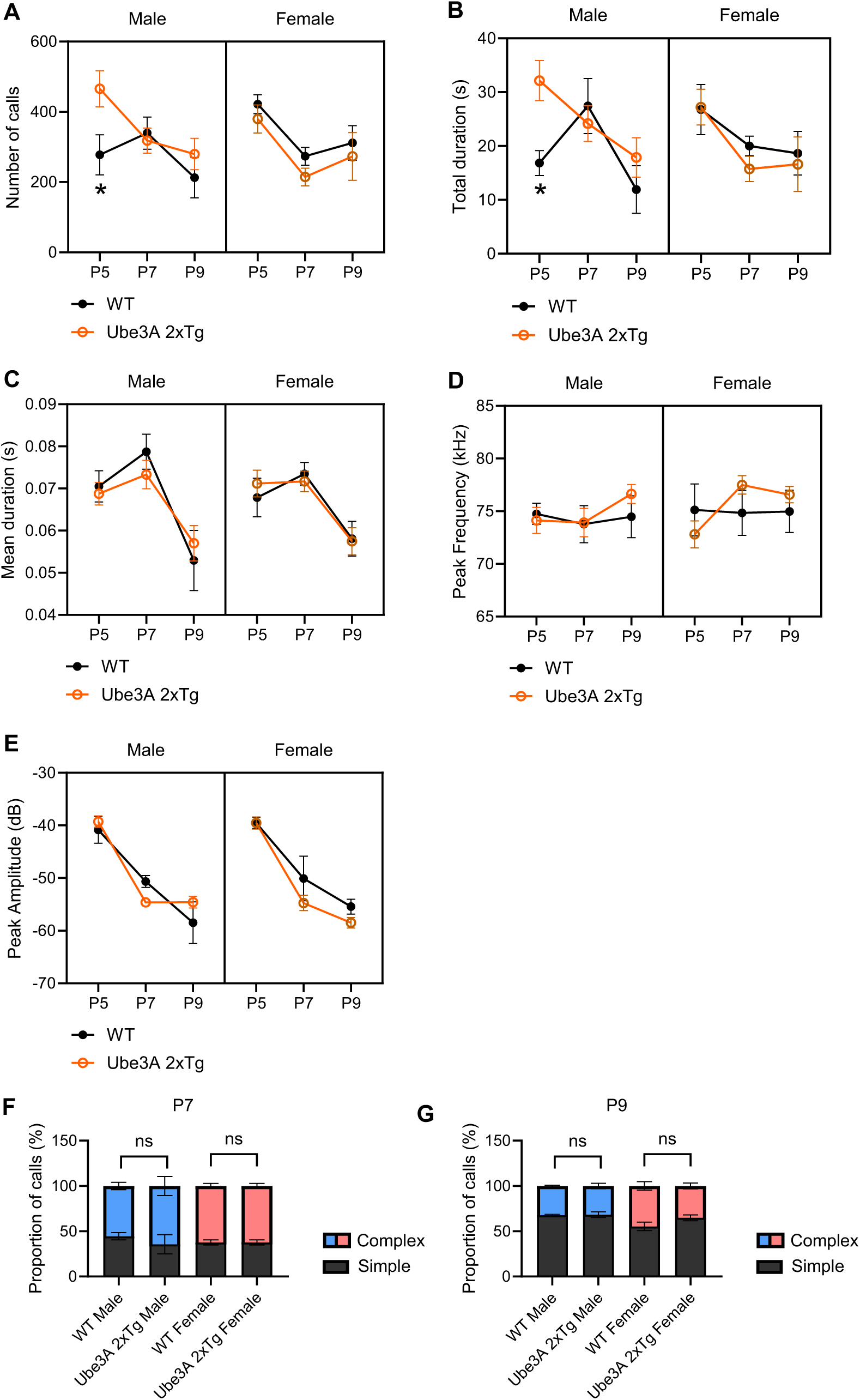
Time-dependent changes of USVs in *Ube3A* 2xTg mice. **(A-E)** Quantifications of the number of calls (A), total call duration (B), mean call syllable duration (C), peak frequency (D), and peak amplitude (E) for USVs recorded on P5, P7, and P9. *Ube3A* 2xTg male mice displayed higher number of calls and longer total call duration than WT males only at P5. No change was observed in *Ube3A* 2xTg females. **(F-G)** Both male and female *Ube3A* 2xTg animals exhibited call syllable types comparable to those of their WT counterparts at P7 (F) and P9 (G). Mean ± SEM. *p<0.05. Three-way ANOVA with Bonferroni’s multiple comparisons test.

## Notes

### Competing Interest Statement

The authors have declared no competing interest.

